# Modeling the schizophrenias: subunit-specific NMDAR antagonism dissociates frontal *δ* and hippocampal *θ* modulation of ~140 Hz oscillations

**DOI:** 10.1101/191882

**Authors:** Benjamin Pittman-Polletta, Kun Hu, Bernat Kocsis

## Abstract

NMDAR antagonism alters mesolimbic, hippocampal, and cortical function, acutely reproducing the positive, cognitive, and negative symptoms of schizophrenia. These physiological and behavioral effects may depend differentially on NMDAR subtype-and region-specific effects. The dramatic electrophysiological signatures of NMDAR blockade in rodents include potentiated high frequency oscillations (HFOs, ~140 Hz), likely generated in mesolimbic structures, and increased HFO phase-amplitude coupling (PAC), a phenomenon related to goal-directed behavior and dopaminergic tone. This study examined the impact of subtype-specific NMDAR antagonism on HFOs and PAC. We found that positive-symptom-associated NR2A-preferring antagonism (NVP-AAM077), but not NR2B-specific antagonism (Ro25-6985) or saline control, replicated increases in HFO power seen with nonspecific antagonism (MK-801). However, PAC following NR2A-preferring antagonism was distinct from all other conditions. While *θ*-HFO PAC was prominent or potentiated in other conditions, NVP-AAM077 increased *δ*-HFO PAC and decreased *θ*-HFO PAC. Furthermore, active wake epochs exhibiting narrowband frontal *δ* oscillations, and not broadband sleep-associated *δ*, selectively exhibited *δ*-HFO coupling, while paradoxical sleep epochs having a high CA1 *θ* to frontal *δ* ratio selectively exhibited *θ*-HFO coupling. Our results suggest: (1) NR2A-preferring antagonism induces oscillopathies reflecting frontal hyperfunction and hippocampal hypofunction; and (2) HFO PAC indexes cortical vs. hippocampal control of mesolimbic circuits.

## Introduction

NMDA hypofunction is one of the leading hypotheses of schizophrenic pathophysiology^1, 2^, mainly due to the ability of NMDA receptor (NMDAR) antagonists such as ketamine and phencyclidine to acutely reproduce the positive, cognitive, and negative symptoms of schizophrenia. NMDARs play a variety of roles in different brain structures, making it challenging to understand how NMDA hypofunction impacts different cells, circuits, and systems to bring about its multiple behavioral effects. Indeed, the recent genetic delineation of multiple schizophrenia subtypes^3^ suggests that NMDA hypofunction – in its replication of multiple symptom types – may actually model several “schizophrenias” simultaneously. Critical details are missing, however. For example, while several experiments have examined the effects of ketamine on fast and slow oscillations and their phase coupling^4–7^, it remains unknown how these changes depend on ketamine’s multiple pharmacological actions, including actions on NMDARs, D2 dopamine (DA) receptors, and HCN1 channels^8–10^.

A natural way to link molecular changes, such as NMDA hypofunction, to behavioral outcomes, such as schizophrenic symptoms, is to study mesoscale phenomena – signatures of coordinated population activity such as brain rhythms in the electroencephalogram (EEG) and local field potentials (LFP)^11^. Both schizophrenia and NMDAR antagonist drugs induce an array of changes in brain rhythms and their coordination^11–13^. Recently, the study of brain rhythms has provided support for the hypothesis that the multiplicity of effects of NMDAR antagonism results in part from the heterogeneity of NMDAR subtypes. Antagonism of the NR2A receptor subtype, but not antagonism of NR2B, C or D receptor subtypes, has been shown to replicate the changes in *γ* power seen with positive symptoms in schizophrenia and with non-specific NMDAR antagonists^14^. Thus, NR2A receptor hypofunction may play a key role in mediating schizophrenia’s positive symptoms.

One of the most marked and robust oscillatory changes observed with acute NMDAR antagonism in rodents is the potentiation of so-called high-frequency oscillations (HFOs, ~120–160 Hz)^4–7^. These narrowband, highly coherent rhythms are carried by mesolimbic circuits, modulating neuronal firing in the VTA^15^ and amygdala^16, 17^ as well as current sources and sinks in the nucleus accumbens (NAcc)^18, 19^. HFO amplitude is often modulated by the phase of low-frequency rhythms in the *θ* (~6–11 Hz) and *δ* (~1–4 Hz) bands. Notably, the mesolimbic *θ and δ oscillations that modulate HFO amplitude are driven by rhythmic activity generated in hippocampus*^20, 21^ and prefrontal cortex (PFC)^22–24^, respectively^15, 17^. Evidence indicates that inputs from PFC and hippocampus compete for control over mesolimbic circuits^25–33^. If HFOs predominantly reflect mesolimbic population activity^15–19^, their phase modulation may reflect the strength of frequency-specific inputs to mesolimbic circuits, either from cortex at *δ* frequencies or from hippocampus at *θ* frequencies.

While several experiments have examined the effects of ketamine on HFO power and its phase coupling to low frequency oscillations^4–7^, fewer studies have used an NMDAR blocker with higher specificity to determine how increases in HFO power depend on ketamine’s multiple pharmacological effects^8, 9, 19^, and none have examined the possible contributions of NMDAR subtype-specific antagonism to altered HFO activity. Nor, to our knowledge, has the relationship between HFO PAC and cortical and hippocampal oscillations been explored directly. To fill this gap, we examined the effects on rhythmic activity and PAC of three NMDAR antagonists previously shown to differentially affect *γ* oscillations^14, 34^: nonspecific antagonist MK-801, NR2A-preferring antagonist NVP-AAM077, and NR2B-specific antagonist Ro25-6985. Our experiments allowed us to observe the coordination between simultaneous changes in HFO power and frontal and hippocampal oscillatory population activity, using simultaneous recordings from frontal cortex, occipital cortex, and hippocampal CA1, while taking advantage of the fact that HFOs are volume conducted from mesolimbic circuits to other brain regions including hippocampus^18^ and frontal cortex^19^.

We found that MK-801 and NVP-AAM077, in contrast to their comparable effects on *γ* activity^14^ and HFO power, induced markedly different patterns of HFO PAC: while MK-801 potentiated *θ*-HFO PAC, NVP-AAM077 potentiated *δ*-HFO PAC and decreased *θ*-HFO PAC. This striking divergence was evident even when examining only active waking (AW) or quiet waking/deep sleep (QW/nREM) epochs, and was paralleled by a dramatic decrease in CA1 *θ* power following NVP-AAM077 injection. Motivated by findings that a narrowband *δ* rhythm – distinct from the broadband *δ* seen during sleep and inactivity – occurs during waking^23, 24^, we segregated AW epochs containing narrowband frontal *δ* in all of our recordings, and found that *δ*-HFO PAC was selectively associated with narrowband frontal *δ* as well as clearly discernable, if rare, following all injections. Similarly, rapid eye movement sleep (REM) epochs with a high CA1 *θ*/frontal *δ* ratio selectively exhibited high levels of *θ*-HFO PAC in all four conditions. Finally, the magnitudes of *θ*-and *δ*-HFO coupling were synchronized with CA1 *θ* and frontal *δ* power, even when examining only AW epochs.

Our findings provide evidence for a link between frontal *δ* oscillations, CA1 *θ* oscillations, and HFO phase-coupling. They suggest that NR2A-preferring antagonism may potentiate *δ* oscillations in frontal cortex and inhibit hippocampal *θ* rhythm, changing the balance of mesolimbic information processing between frontal and hippocampal afferents. These results also suggest that blockade of different NMDAR subtypes can have markedly different effects on the mesoscale physiology of the brain, perhaps resulting in receptor subtype-specific behavioral alterations that may correspond to schizophrenia symptom subtypes and explain the multiple behavioral effects of nonspecific NMDAR antagonism.

## 1 Methods

### 1.1 Recordings

All experiments were performed in accordance with National Institute of Health guidelines and were approved by the Institutional Animal Care and Use Committee of Beth Israel Deaconess Medical Center. Six (*n* = 6) rats were housed in a temperature and humidity-controlled room with 12h/12h light/dark cycle; food and water was available ad libitum both in the home cage and during recordings. Stainless steel screws were used to record EEG over the frontal (1 mm anterior and 2 mm lateral to bregma) and occipital (6.5 mm posterior and 3 mm lateral to the bregma) cortices, and pairs of twisted wires were implanted in the hippocampus (CA1, as identified by the phase of *θ* relative to occipital cortex) to record LFPs. Two additional screw electrodes were inserted ~5 mm anterior to bregma and over the cerebellum for ground and reference, and electromyograms (EMG) were recorded using multithreaded wires in the neck muscles. Recordings began after a 7-to 10-day recovery period following chronic implantantion, and experiments with drug injections started after several days of control recordings. Each rat was injected (in 1 mL/kg volume, subcutaneously, on different days and with at least 3 days in between) with nonselective NMDAR antagonist MK-801 (0.2 mg/kg, Tocris), NR2A-preferring antagonist NVP-AAM077 (20 mg/kg, Novartis), NR2B-selective antagonist Ro25-6985 (10 mg s/c, Tocris), and saline vehicle as a control. Recordings started in the early morning, beginning ≥ 3 h prior to injection and ending ≥ 16 h after injection. Data were divided into ~16 s epochs for further analysis.

### 1.2 Spectral Analyses

Power spectra were calculated for each epoch using the Thompson multitaper method. Power at each frequency was expressed as a percentage of mean baseline power, defined to be the mean power (at a given frequency) over the 4 h preceding injection. Power within the *δ* (1–5 Hz), *θ* (5–11 Hz), high *γ* (40–90 Hz), and HFO (120–160 Hz) bands was calculated by summing spectral power over the frequencies within these bands.

### 1.3 Vigilance States

EMG power and CA1 *θ* power were used to automatically categorize each epoch into one of three behavioral states: active waking (AW), characterized by high *θ* and high EMG power; paradoxical sleep (REM), characterized by high *θ* power and low EMG power; and quiet waking/non-REM sleep (QW/nREM), characterized by low *θ* and low EMG power. Candidate REM epochs were further checked by manual scoring.

### 1.4 PAC Analyses

PAC comodulograms were computed for all epochs. First, oscillations were extracted from each epoch via convolution with 7-cycle complex Morlet wavelets, having central frequencies spaced every 0.25 Hz from 1 to 12 Hz and every 5 Hz from 20 to 200 Hz. Instantaneous amplitude and phase time series *A_f_* (*t*) and *φ_f_* (*t*) were calculated for each frequency *f* as the absolute value and angle of the resulting complex time series. For each pair of high-frequency (20-200 Hz) amplitude and low-frequency (1–12 Hz) phase time series, an inverse entropy index^35^ was computed. The resulting inverse entropy measure (IE) quantifies the degree of phase-amplitude dependence. An IE of zero indicates no phase-amplitude dependence (a uniform distribution of amplitude with respect to phase). An IE of one is obtained when all the amplitude occurs in a single phase bin. In general, higher IE is awarded to distributions exhibiting a stronger dependence of amplitude on phase, including distributions exhibiting multiple peaks.

For a finite data series, IE will always take finite nonzero values, even when no coupling is present in the data. To remove these non-coupling related influences, we used surrogate data to estimate “background” values of IE. These background IE values were estimated using epoch-shuffled surrogate data: for each h relative to injection in each subject, and for each frequency pair, 1000 random pairings of non-simultaneous high-frequency amplitude and low-frequency phase time series were used to calculate a distribution of surrogate IE values. For each pair of frequencies, this distribution was used to z-score the observed IE, yielding a z-score of coupling significance or modulation index (MI) for that epoch and pair of frequencies.

### 1.5 Isolating Epochs of Narrowband *δ*

To determine epochs containing narrowband *δ*, we calculated the entropy of the spectral power in the *δ* band. The spectrum obtained using the Thompson multitaper method was smoothed by convolution with a Gaussian of standard deviation 0.075 Hz. The smoothed spectrum within the *δ* range (0.5 – 4.5 Hz) was normalized to have unit sum. The entropy of this distribution of *δ* power by frequency was computed, resulting in a measure of the uniformity of the distribution of spectral power across frequencies (*δ* entropy). The higher the *δ* entropy value, the more uniformly power is distributed across the *δ* band; the lower the *δ* entropy value, the “peakier” is the distribution of spectral power in the *δ* band. Next, we calculated the values of *δ* entropy corresponding to the first and last percentile of observed *δ* entropy across all recorded waking epochs, for each drug. Epochs having a *δ* entropy less than or equal to the first percentile were considered to exhibit narrowband *δ*, while epochs having a *δ* entropy greater than or equal to the last percentile were considered to exhibit broadband *δ*. Median PAC comodulograms were plotted for these two subsets of epochs.

### 1.6 Correlating Spectral Power and PAC

To correlate spectral power and PAC, values of MI for all frequency pairs, and values of power in the *θ* and *δ* bands, were correlated during the first 4 h following injection in all conditions.

### 1.7 Isolating Epochs with High *θ*/*δ* Ratio

To determine epochs containing high *θ*/*δ* ratio, we took the ratio of the band power within the *θ* and *δ* bands. Within each vigilance state, the epochs having *θ*/*δ* ratio within the first and last percentile of observed *θ*/*δ* ratios was selected, by the same procedure as for *δ* entropy, above. Median PAC comodulograms were plotted for these low and high *θ*/*δ* ratio epochs.

### 1.8 Statistics

As IE and spectral power take positive values, they are not normally distributed (even after z-scoring against surrogate distributions). This was confirmed using Kolmogorov-Smirnov tests. Thus, median values are used in visualizations, and ranksum tests were used for comparisons to saline. To quantify the similarities and differences in IE and power between drugs pre-and post-injection, we calculated pairwise ranksum tests (comparing each drug to saline injection) for each six min period relative to injection. These tests compared the *n* = 6 observations from each rat (averaged over the six min period) for drug and saline. Tests were performed on sums of MI over frequency ranges of interest and on baseline-normalized band power.

## 2 Results

### 2.1 MK-801 and NVP-AAM077 Have Similar Effects on High Frequency Rhythms

Both MK-801 and NVP-AAM077 administration were observed to result in prolonged periods (~4–6 hs) of wakefulness, as opposed to Ro25-6985 and saline. The animals mostly exhibited abnormal “stereotyped” behaviors, such as head-banging, prolonged climbing-like movements in the corner of the cage, etc.^14, 34^ During these four hs, high *γ* and HFO power increased dramatically in these two conditions (Fig. 1), roughly following the time course of the abnormal low *γ* increases reported following these drugs^14, 36, 37^. In contrast, Ro25-6985 did not induce persistent changes in fast oscillations^34^.

**Figure 1.**
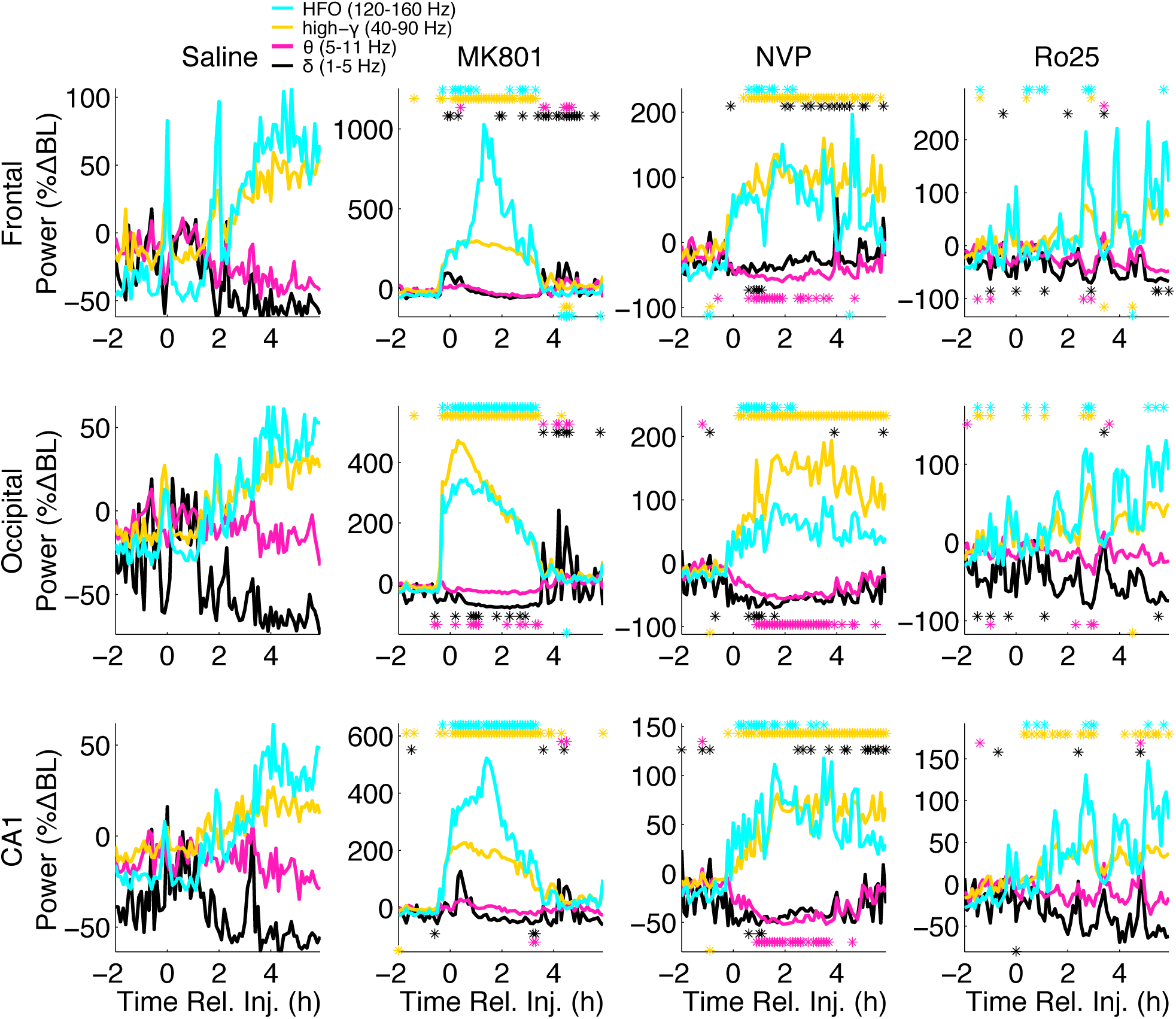
NVP-AAM077 and MK-801 induce similar increases in high frequency power, but have different effects on low frequency power. Median (n=6) spectral power (normalized as a percentage change from mean baseline power) summed over the *δ* (1–5 Hz), *θ* (5–11 Hz), high *γ* (40–90 Hz), and HFO (120–160 Hz) bands are shown for frontal (top), occipital (middle), and CA1 (bottom) electrodes, following saline (left), MK-801 (middle left), NVP-AAM077 (middle right), and Ro25-6985 (right) injection. Stars indicate 6 min periods for which spectral power is significantly higher (above) or lower (below) than saline (ranksum test, significance level = 0.05).

MK-801 and NVP-AAM077 (but not Ro25-6985) also affected slow oscillations. Previous studies^6, 37^ reported layer dependent changes in hippocampal *θ* power after MK-801 administration, with *θ* decreases in superficial layers of CA1 accompanying strong *θ* increases in deep electrodes placed at or below the hippocampal fissure. In our current sample, *θ* showed little change following MK-801 administration, but was strongly suppressed (relative to saline) after NVP-AAM077 administration. This suppression lasted for 4 h in occipital cortex and CA1, and for 2 h in frontal cortical recordings. Following NVP-AAM077, power in the *δ* band showed a biphasic reaction: it was suppressed in the first h, and then started to rise and reached significantly elevated levels in the frontal cortex during the 3*^rd^* to 6*^th^* h post-injection (Fig. 1, right middle column). In contrast, *δ* did not increase following MK-801until 4 h after injection, when changes in HFO and high *γ* (and PAC) had worn off.

### 2.2 MK-801 and NVP-AAM077 Induce Primarily *θ*-HFO and *δ*-HFO PAC, Respectively

Following saline injection (Figs. 2, S1), we observed region-specific profiles of PAC. In frontal cortex, the most prominent coupling was between *δ* phase and high *γ* power, with weaker coupling between *θ* and high *γ*/HFOs. In occipital cortex and CA1, prominent *θ*-HFO PAC was observed. In CA1, weak *θ*-*γ* PAC was also visible.

**Figure 2.**
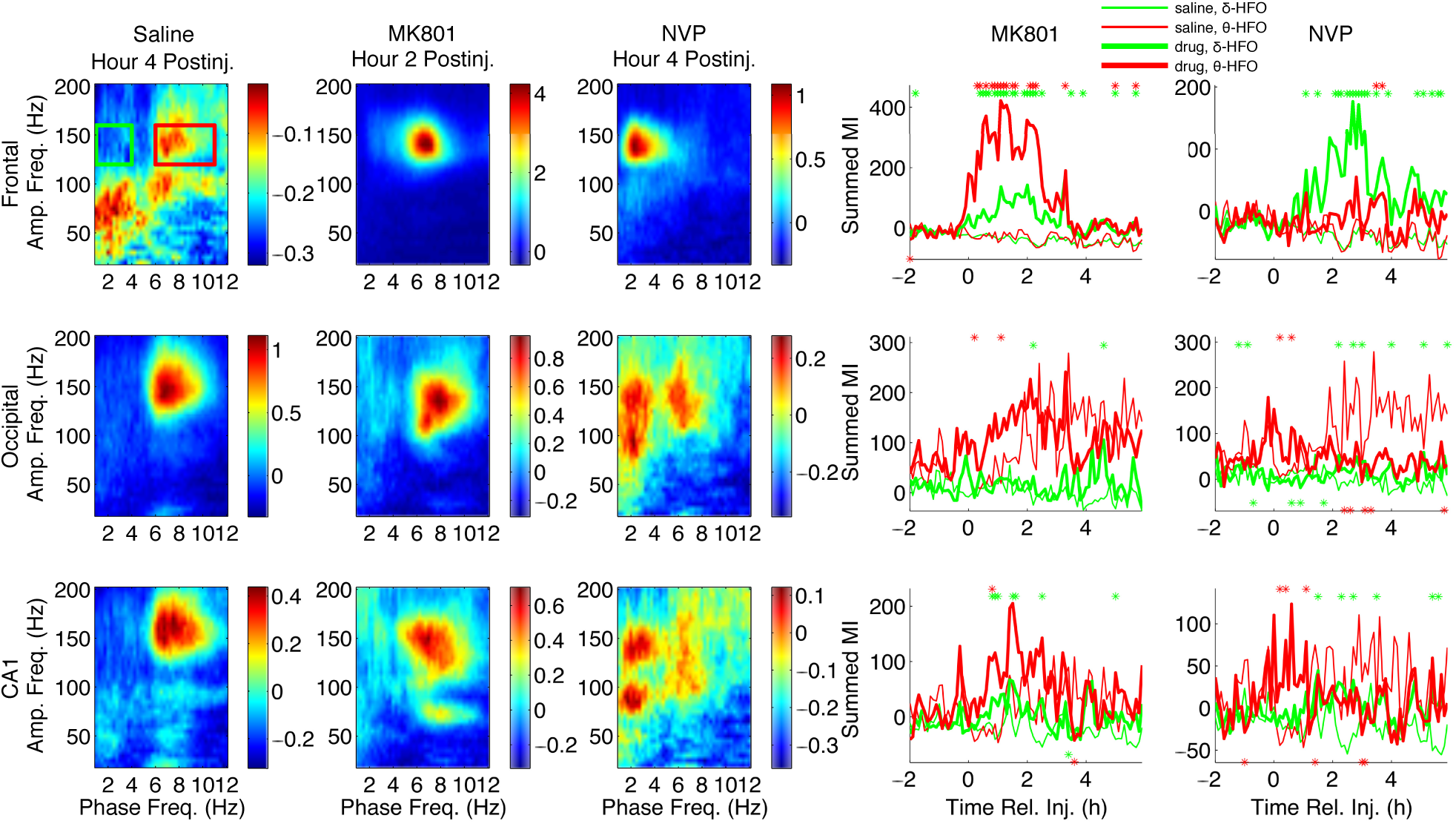
Patterns of phase-amplitude coupling following MK-801 and NVP-AAM077 injection differ markedly. In the left three columns, median (n=6) comodulograms for the h of largest effect are shown for simultaneous LFP recordings from frontal (top row), occipital (middle row), and CA1 (bottom row) electrodes, following injection of saline, MK-801, and NVP-AAM07. These comodulograms are also shown in Figures 3, 4, and S1; here they are shown with individually optimized color scales, to more clearly show the pattern of PAC in each region during the h of largest effect for each drug. The right two columns show timeseries of summed PAC over the *θ*-HFO (red rectangle and time series) and *δ*-HFO (green rectangle and time series) frequency ranges from 2 h pre-injection to 6 h post-injection. Thick lines are drug time series; thin lines are saline time series; stars indicate 6 min periods for which drug PAC is significantly higher (above) or lower (below) than saline (ranksum test, significance level = 0.05).

In agreement with previous reports for ketamine^5–7^, median PAC comodulograms following MK-801 injection revealed dramatic increases in *θ*-HFO PAC in all brain regions relative to saline injection (Figs. 2, 3), suggesting that ketamine increases PAC^5–7^ through its action at NMDARs, and that its D2 receptor agonism is not necessary for increased coupling. PAC increases were most dramatic in frontal cortex, followed by occipital cortex, and then by CA1. These increases in *θ*-HFO PAC were accompanied by statistically significant but less dramatic increases in *δ*-HFO PAC. In CA1, MK-801 injection also led to an increase in *θ*-*γ* PAC (Figs. 2, 3).

**Figure 3.**
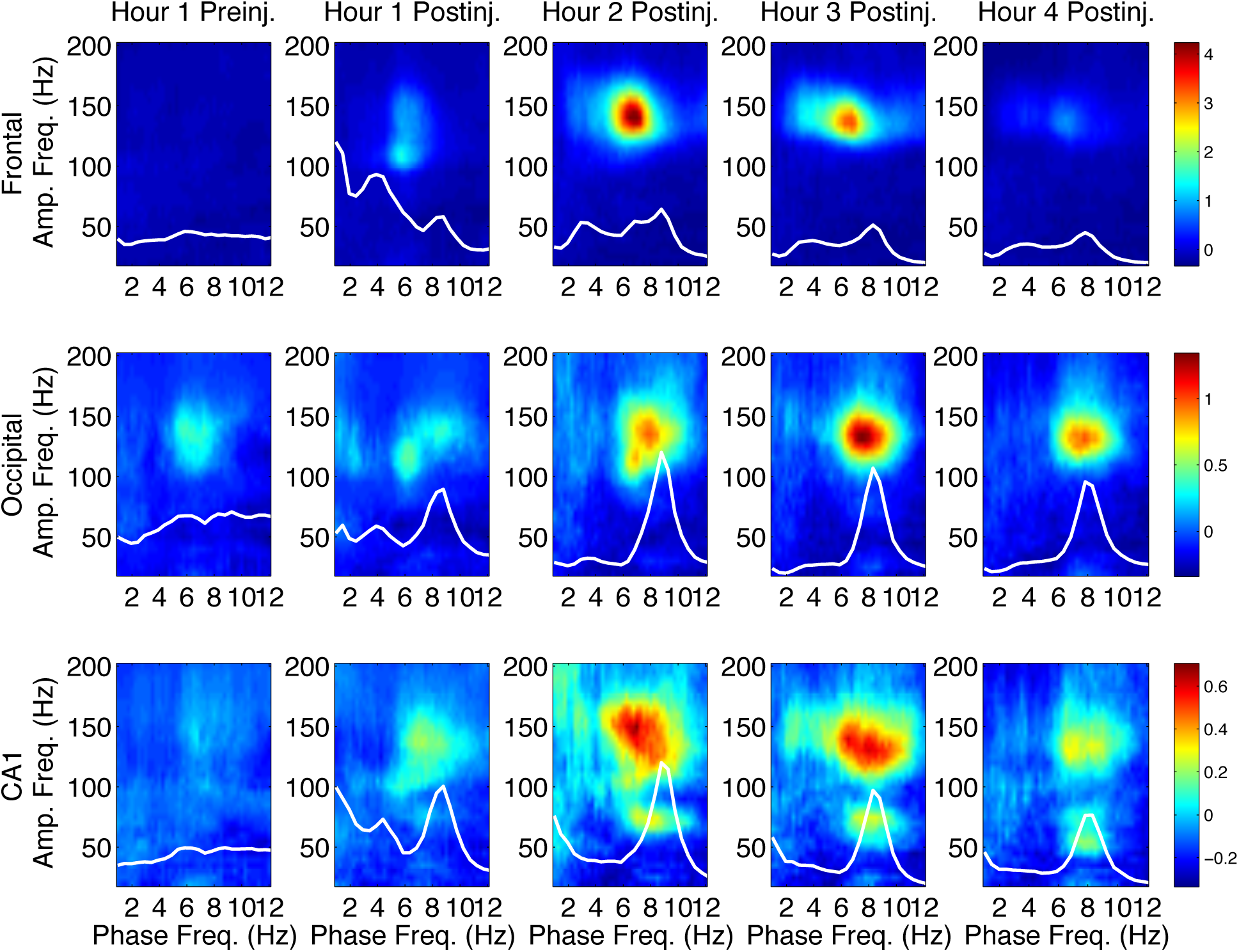
MK-801 increases *θ*-HFO PAC. Median (n=6) comodulograms for the 1 h preceding and the 4 h following MK-801 injection, shown for frontal (top), occipital (middle), and CA1 (bottom) electrodes (simultaneous recordings). Overlaid white traces show the mean spectral power at frequencies below 12 Hz for each h.

Following NVP-AAM077 injection, the largest observed effect was an increase in *δ*-HFO PAC at all sites, relative to saline (Figs. 2, 4). In contrast, only minor increases in *θ*-HFO PAC were seen in frontal cortex, and decreases in *θ*-HFO PAC were observed in occipital cortex and CA1, relative to saline (Fig. 2, rightmost column, Fig. 4). No changes in *θ*-*γ* or *δ*-*γ* PAC were observed. This divergence is especially noteworthy, given that NVP-AAM077 is only NR2A-preferring (not strongly NR2A-selective). Following injection of Ro25-6985, few prolonged changes in PAC were observed (not shown).

**Figure 4.**
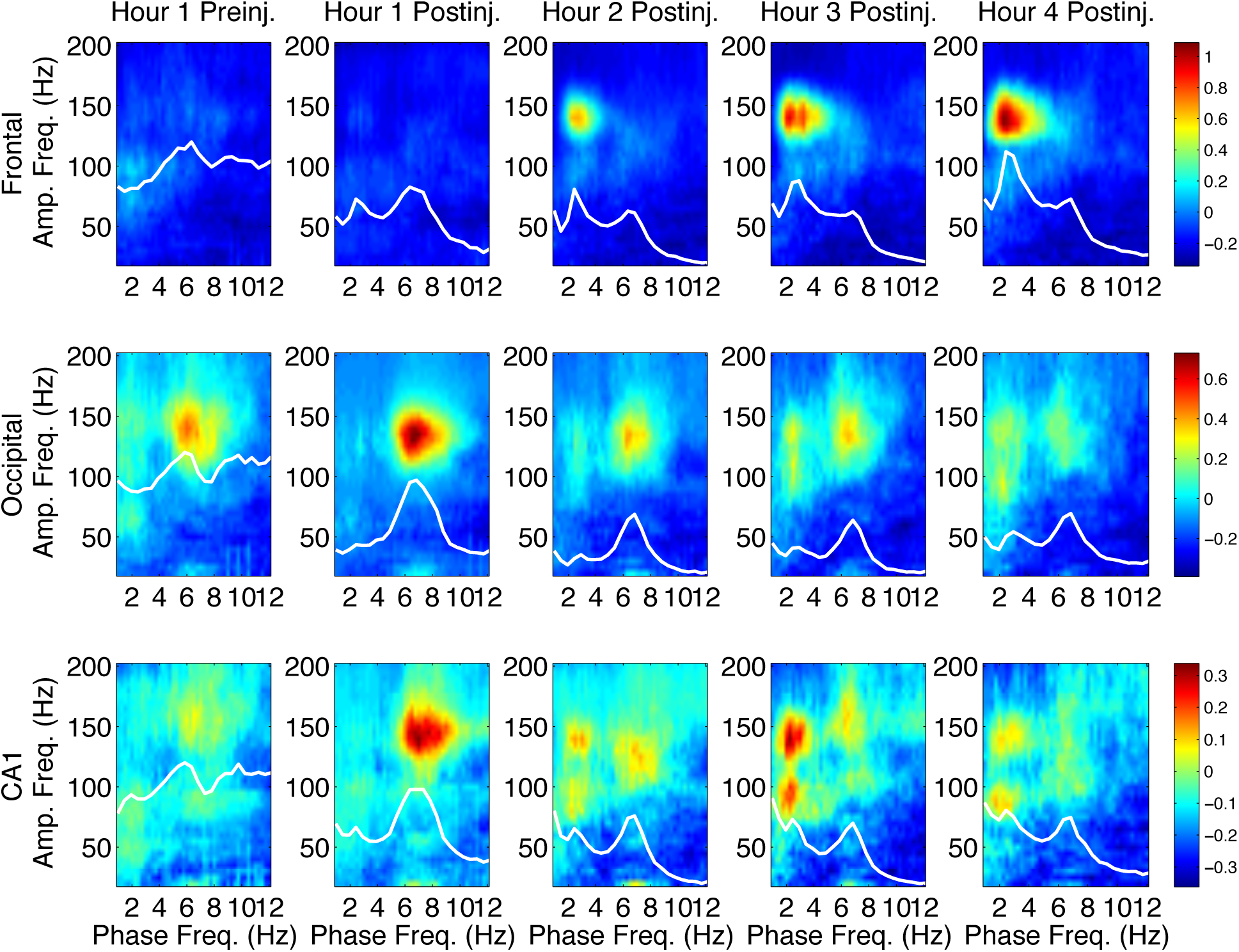
NVP increases *θ*-HFO PAC. Median (n=6) comodulograms for the 1 h preceding and the 4 h following NVPAAM077 injection, shown for frontal (top), occipital (middle), and CA1 (bottom) electrodes (simultaneous recordings). Overlaid white traces show the mean spectral power at frequencies below 12 Hz for each h.

*θ* coupling of *γ* and HFO power is known to be higher during *θ* states such as AW and REM. As vigilance states are usually defined by EMG activity and EEG *θ* power (see Section 1.3), behavioral analyses of our data using traditional terms and scoring may be confounded by the decreases in EEG *θ* power observed following NVP-AAM077 injection. Rats were awake 4–6 h after both MK-801 and NVP-AAM077 injections^14, 34^. Following MK-801, even though the animals were not engaged in normal *θ*-associated exploratory behavior, increased EMG activity was mostly accompanied by *θ*, and an increase in traditionally scored active wake (AW) epochs relative to saline was observed (Fig. S2). On the other hand, there were fewer *θ* states following NVP-AAM077 injection, so observed increases in wakefulness following NVP-AAM077 were not reflected in a change in the proportion of AW epochs (Fig. S2), but could be reflected in an increase in traditionally scored quiet wake (QW) epochs and a simultaneous decrease in traditionally scored non-REM sleep (nREM) epochs relative to saline. Thus, to address the relationship between changes in vigilance states and changes in HFO PAC, in addition to behavioral analyses using the traditional vigilance state categories (Fig. S2, see Section 1.3), we also restricted our analyses to only AW epochs (Fig. S3), or only QW/nREM epochs (Fig. S4), and found that the distinct patterns of PAC described above persisted. Thus, MK-801 induced increased *θ*-HFO coupling, relative to saline, within both AW and QW/nREM epochs; and NVP-AAM077 induced increased *δ*-HFO coupling and decreased *θ*-HFO coupling, relative to saline, within both AW and QW/nREM epochs.

Evidence suggests that NMDAR antagonist-potentiated HFOs have current generators in mesolimbic circuits, and are then volume conducted to other brain regions^15–19^. However, spontaneous HFOs may be generated locally in other brain regions, including hippocampus^18^. We observed small differences in the time course and spectral parameters of HFO power between recording sites, and differences between sites in HFO PAC, indicating that the predominance of *δ* and *θ* oscillations in frontal cortex and CA1, respectively, might bias the detection of *δ*-HFO PAC in frontal cortex and the detection of *θ*-HFO PAC in CA1. Thus, for further analyses of the precise timecourse and dynamics of HFO phase modulation, and its correlation with *δ* and *θ* power, we utilized only measurements of PAC taken from the “neutral” occipital cortex.

### 2.3 Following NVP-AAM077 Injection, *δ*-and *θ*-HFO Phase Coupling are Dissociable in Time

To explore the relationship between *θ*-HFO and *δ*-HFO PAC appearing in occipital cortex following NVP-AAM077 injection on a short time scale, we examined individual epochs and 6 min segments of EEG. While both *θ*-HFO and *δ*-HFO PAC were observed in occipital cortex during the h of greatest effect following NVP-AAM077 injection (Fig. 5, top left), the top quartiles of epochs for *θ*-HFO and *δ*-HFO PAC during this h (Fig. 5, top middle and right) revealed a dissociation between these two phenomena. In individual animals, the time-courses of *δ*-HFO and *θ*-HFO PAC at 6 min resolution were not aligned (Fig. 5, bottom left), and in the 4 h following injection, the correlation between *δ*-HFO PAC and *θ*-HFO PAC across animals was not significant (p = 1), with the two variables showing a slightly negative relationship (Fig. 5, bottom right).

**Figure 5.**
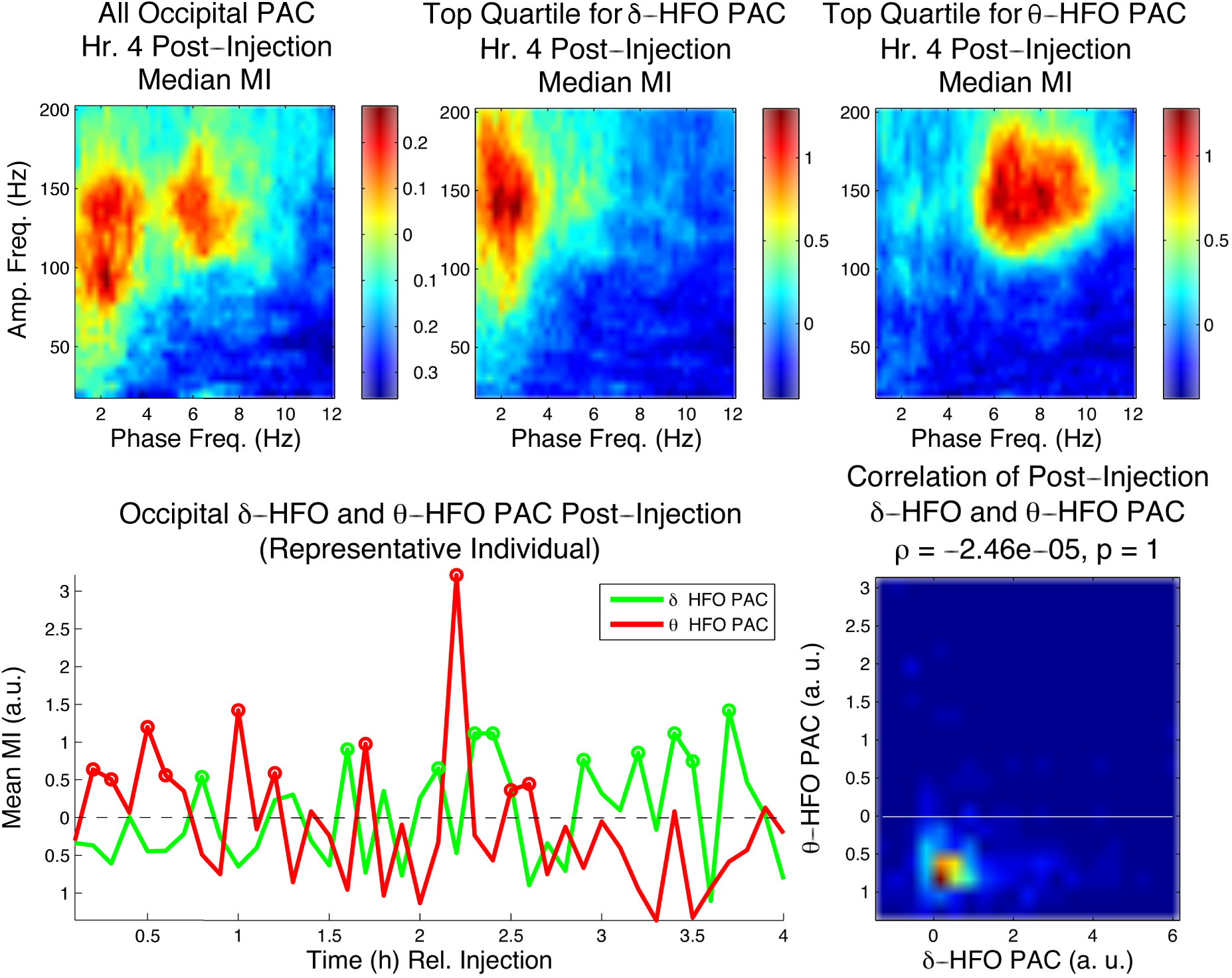
*δ*-HFO PAC and *θ*-HFO PAC appear at different times in occipital cortex following NVP-AAM077 administration. Top left, the profile of median (n=6) occipital PAC in the 4*^th^* h following NVP-AAM077 injection. Top center and right, the profiles of median (n=6) PAC for the top quartile of epochs with respect to *δ*-HFO and *θ*-HFO PAC. Bottom left, time courses of summed *δ*-HFO and *θ-*HFO PAC for 6 min periods following injection (bottom left, representative animal, circles indicate 6 min periods in the top quartile). Bottom right, the correlation between *δ-*HFO PAC and *θ*-HFO PAC over the 4 h following injection for all animals.

In contrast, in the h of greatest effect following MK-801 injection, while the top quartile of epochs for *θ*-HFO PAC exhibited HFO coupling only to *θ* frequencies, the top quartile of epochs for *δ*-HFO PAC exhibited both *δ*-and *θ*-HFO PAC, and *δ*-and *θ*-HFO PAC were significantly correlated (p = 9.17×10^−5^) in the 4 h post-injection (Fig. S5). This suggests that the elevation in *δ*-HFO PAC seen following MK-801 injection was not independent of increased *θ*-HFO PAC.

### 2.4 *δ-*HFO PAC is Restricted to Epochs Containing Narrowband *δ*

Since *δ* power may increase during both sleep and wake, but only wake was seen in the 4 h following NVP-AAM077 injection, we hypothesized that the expression of *δ*-HFO PAC in physiological conditions might depend on the presence of wake-associated narrowband *δ* oscillations^23, 24^. This waking *δ* rhythm is different from the broadband *δ* of inactivity and slow-wave sleep^23, 24^, and may be cognitively important^38^. Thus, we sought to hone in on *δ-*HFO coupling by selecting only those active waking epochs (across all four drugs) in which narrowband *δ* was observed in frontal electrodes.

To establish that *δ* oscillations were narrowband, we calculated the entropy of the spectrum within the *δ* band (treating it as a probability distribution) for the frontal EEG for each epoch (see Methods). The resulting measure of *δ* entropy quantified the uniformity of spectral power across the *δ* band. Thus, high *δ* entropy epochs exhibited broadband *δ*, while low *δ* entropy epochs exhibited “peakier”, narrowband *δ*. Over all waking epochs for each drug, we determined the *δ* entropy values marking the first and last percentiles of observed *δ* entropy values, and used these to define narrowband and broadband *δ* epochs, respectively. Figure 6 compares median comodulograms from occipital electrodes for narrowband and broadband *δ* epochs, for each drug. It shows that *δ*-HFO PAC is exclusively associated with narrowband *δ* epochs, and nearly invisible in broadband *δ* epochs, across conditions.

**Figure 6.**
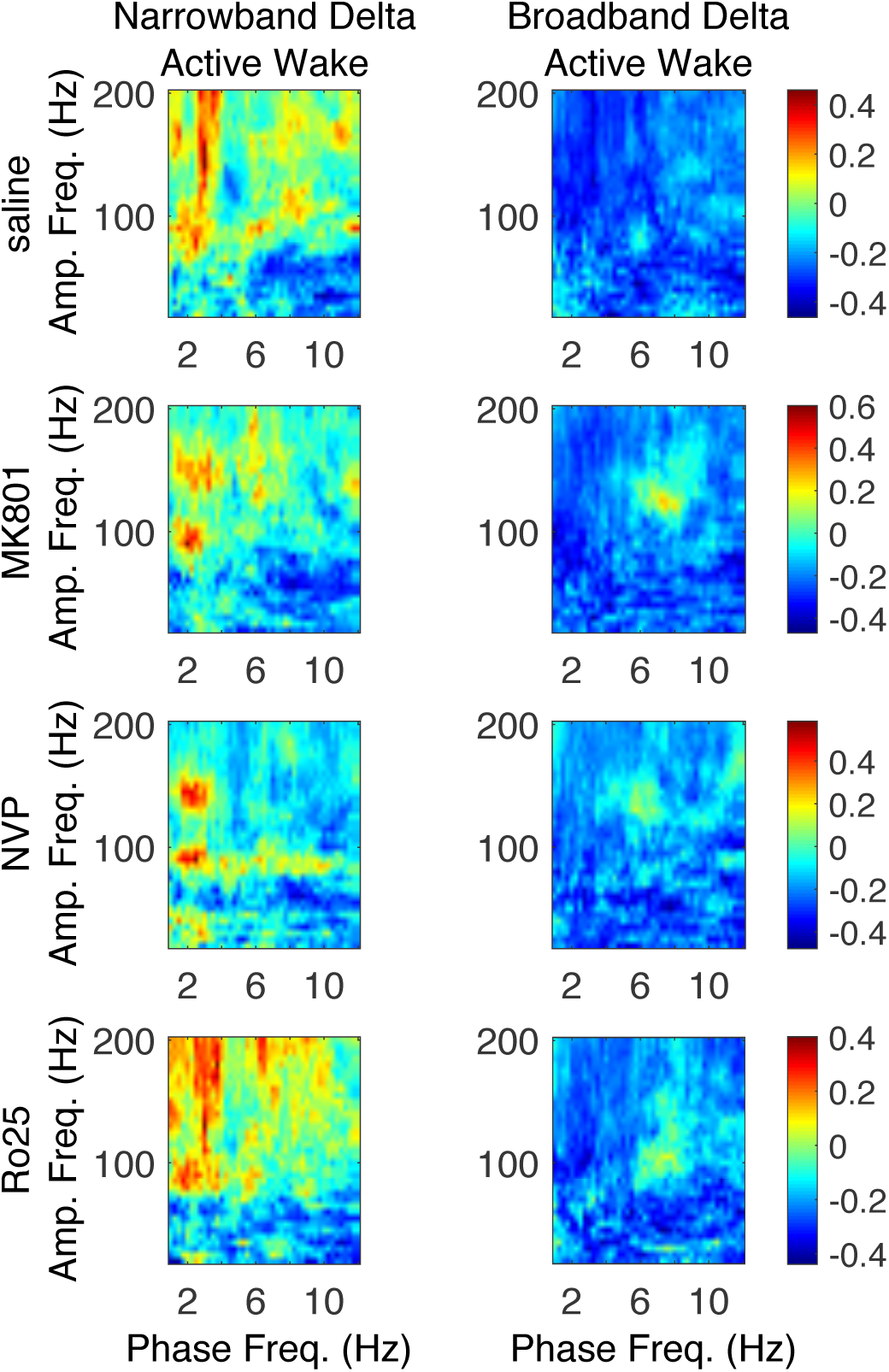
Narrowband frontal *δ* differentiates AW epochs exhibiting *δ-*HFO coupling. Median (n=6) comodulograms for the AW epochs for which the frontal LFP has a narrowband peak in the *δ* band and exhibit *δ-*HFO coupling (left), and for the AW epochs exhibiting broadband frontal *δ* and an absence of *δ-*HFO coupling (right).

### 2.5 *θ*-HFO and *δ*-HFO PAC are Correlated with CA1 *θ* and Frontal *δ*, Respectively

Over the same period that HFO phase-coupling switched from *θ* to *δ* frequencies, spectral analysis revealed a switch in the relative dominance of *θ* and *δ* power in the areas where these slow oscillations most prominently occur and are most likely generated, i.e., in hippocampal CA1 and frontal cortex, respectively^20, 21, 23, 24^ (Fig. 7). In contrast, no switch was seen in the relative dominance of CA1 *δ* and frontal *θ* (not shown). Comparison of baseline-normalized spectral power in the *θ* and *δ* bands between h 1 and h 4 post-injection revealed that while frontal *δ* increased and CA1 *θ* decreased highly significantly (p = 0.0198 and p = 1.173×10^−5^, respectively), frontal *θ* and CA1 *δ* did not change (p = 0.815 and p = 0.931, respectively). These observations held even when we restricted our analysis to AW epochs (Fig. S6; frontal *δ* increase, p = 3.4475×10^−5^; CA1 *θ* decrease, p = 2.1254×10^−4^; frontal *θ*, p = 0.1855; CA1 *δ*, p = 0.1540).

**Figure 7.**
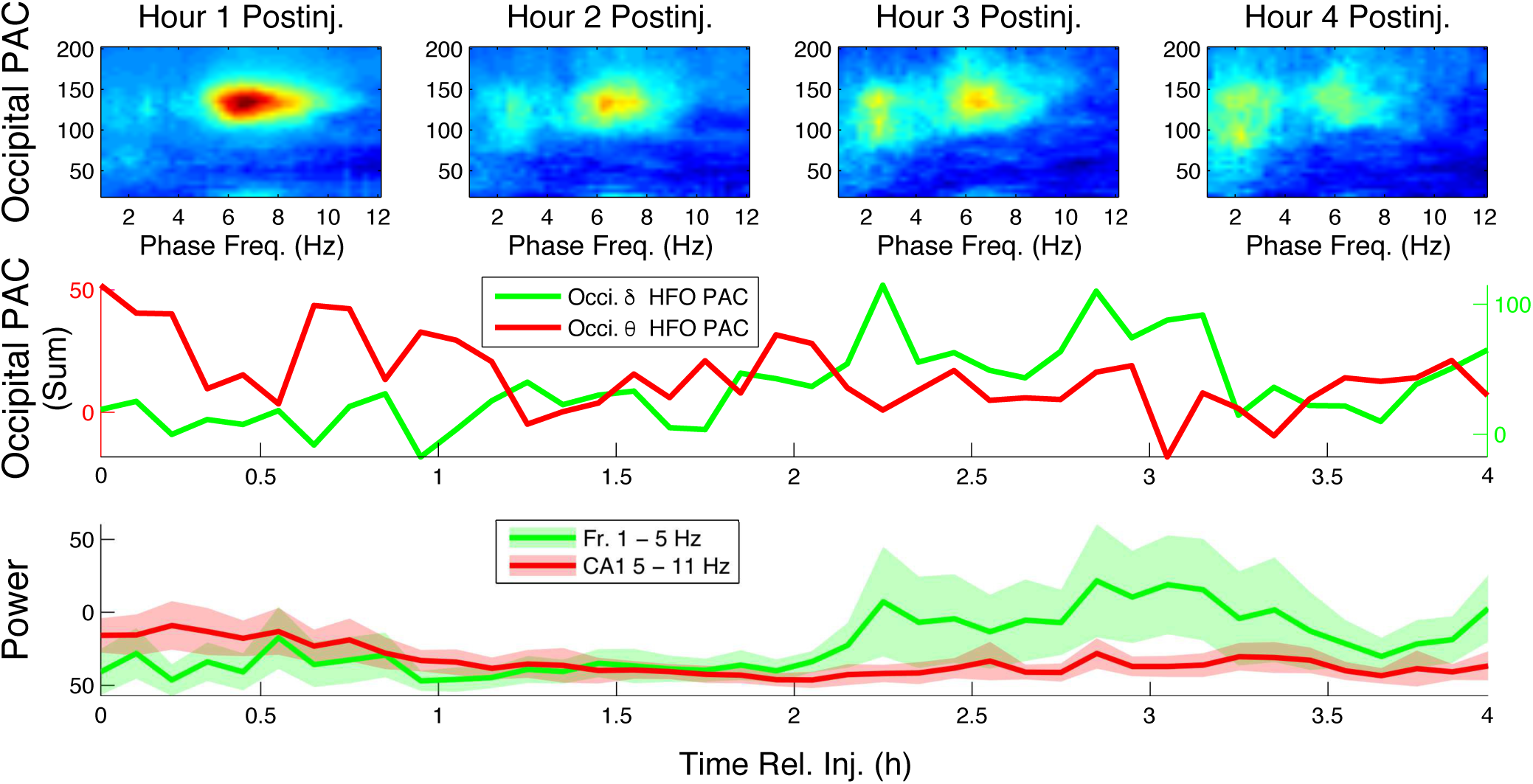
The time courses of frontal *δ* power and CA1 *θ* power mirror the time courses of *δ-*HFO PAC and *θ-*HFO PAC, respectively. Top, median (n=6) occipital comodulograms are shown for the first 4 h following NVP-AAM077 injection. Middle, profiles of median (n=6) summed PAC (at 6 min intervals) are shown for *δ-*HFO and *θ-*HFO ranges. Bottom, mean ± s.d. (n=6) frontal *δ* power (summed over 1–5 Hz) is plotted alongside median (n=6) CA1 *θ* power (summed over 5–11 Hz).

We hypothesized that these synchronous changes in PAC and spectral power were related. To further establish the connection between CA1 *θ* and *θ*-HFO PAC on one hand, and between frontal *δ* and *δ*-HFO PAC on the other, we correlated the values of PAC calculated at all frequency pairs with spectral power summed over the *δ* and *θ* bands in frontal cortex and CA1, respectively (Fig. S7), at the ~16 second timescale of epochs. This analysis was repeated for all injections, including saline and different NMDAR blockers, and revealed a universal relationship between slow oscillations and HFO modulation. The results show in particular that *δ*-HFO and *δ*-high *γ* PAC were positively correlated, whereas occipital and CA1 *θ*-HFO PAC were negatively correlated, with frontal *δ* power (left panels, Fig. S7). In contrast, occipital and CA1 *θ*-HFO PAC were positively correlated with CA1 *θ* power, and *δ*-HFO PAC was negatively correlated with changes in spectral power in the *θ* band in CA1 (right panels, Fig. S7). These observations held for AW epochs alone (Fig. S8). This is especially noteworthy given that *θ* and *δ* power were either slightly positively correlated (following MK-801, *ρ* = 0.194 and p = 2.19×10^−5^; following NVP-AAM077, *ρ* = 0.217 and p = 2.06×10^−6^; following Ro25-6985, *ρ* = 0.111 and p = 0.0152) or uncorrelated (following saline, p = 0.127) in the 4 h post-injection.

### 2.6 *θ*-HFO PAC is Highest During REM Epochs Having High *θ*/*δ* Ratio

To further probe the dependence of *θ*-HFO PAC on hippocampal *θ*, we sought epochs exhibiting high CA1 *θ* and low frontal *δ* power. This oscillatory profile is exhibited during REM, and it has been shown previously that *θ*-HFO PAC is elevated during REM relative to both wake and non-REM sleep^39^. We examined the relationship of NMDAR antagonist potentiated *θ*-HFO PAC to CA1 *θ* oscillations using the *θ* state of REM, and the ratio of CA1 *θ* power to frontal *δ* power. Over all REM epochs for all drugs, we determined the values of the *θ*/*δ* ratio marking the first and last percentiles of observed *θ*/*δ* ratios, and used these to define low and high *θ*/*δ* ratio epochs, respectively. Figure 8 shows that *θ*-HFO PAC is exclusively associated with high *θ*/*δ* ratio REM epochs, and nearly invisible in low *θ*/*δ* ratio REM epochs, across conditions.

**Figure 8.**
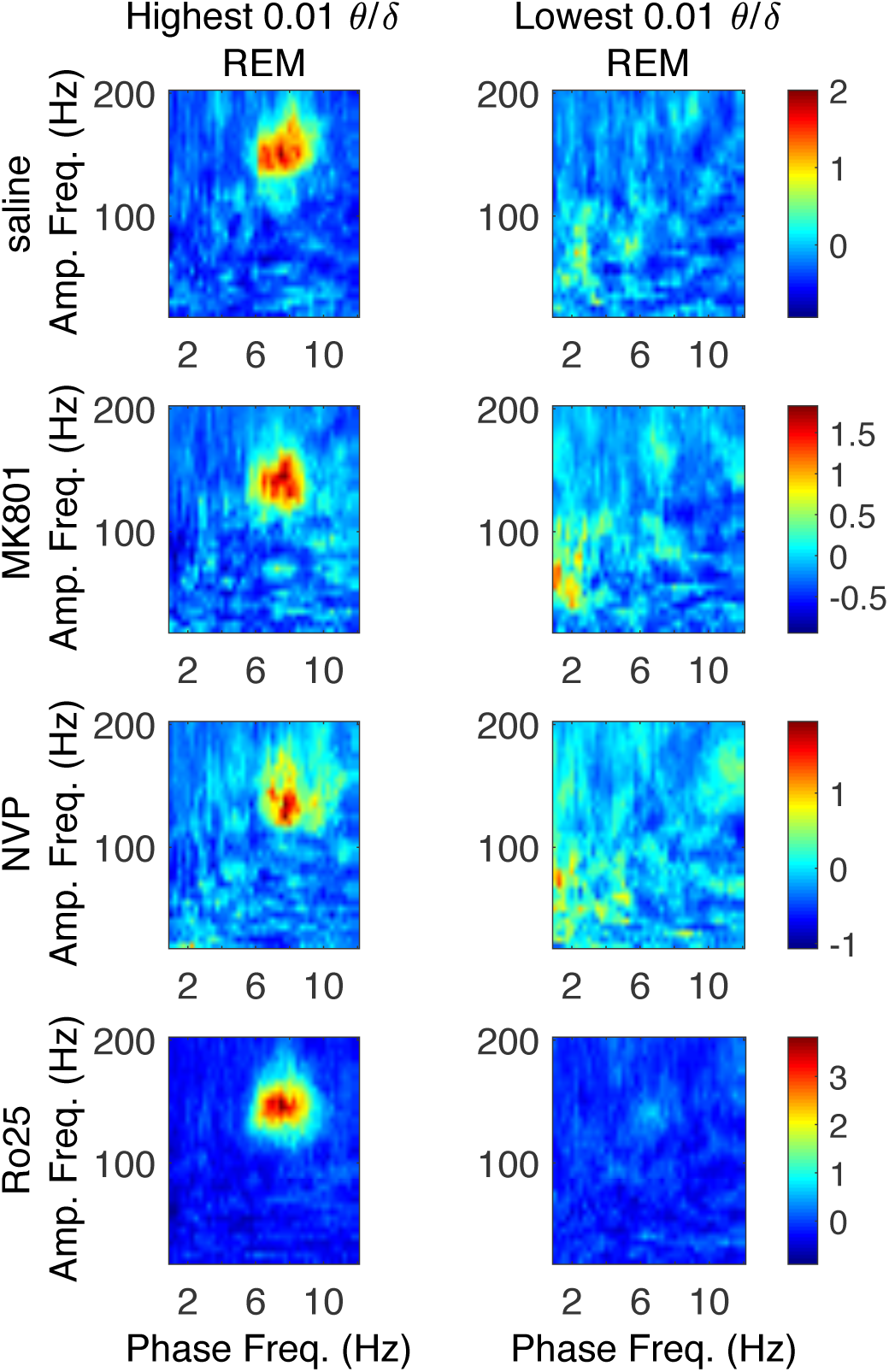
High *θ*/*δ* ratio differentiates REM epochs exhibiting *θ*-HFO coupling. Median (n=6) comodulograms for the REM epochs for which the CA1 LFP has a high *θ*/*δ* ratio (left), and for the REM epochs exhibiting a low *θ*/*δ* ratio (right).

## 3 Discussion

The major findings of this study are that nonspecific NMDAR blockade primarily increased *θ*-HFO coupling shortly after systemic administration of MK-801, whereas the NMDAR subunit-specific NR2A-preferring antagonist NVP-AAM077 induced a delayed increase in *δ*-HFO coupling lasting for h and accompanied by a decrease in *θ*-HFO coupling. These coupling changes did not depend on changes in vigilance state following drug injection. Our observations agree with results on the administration of the non-specific NMDAR antagonist ketamine^4–7, 37^. We show for the first time that changes in HFO phase-coupling following ketamine administration are most likely due to its action on NMDAR and independent of its action at D2 dopamine receptors and HCN1 channels.

Our results allow the formulation of new hypotheses regarding the control and function of mesolimbic circuits. Our observations suggest that subunit-specific changes reflect an association between the dominant frequency of HFO PAC and the power of frontal *δ* and hippocampal *θ* oscillations. Following NVP-AAM077 injection, the remarkable predominance of *δ-*HFO PAC was correlated with frontal *δ* power, and across drugs, the presence of narrowband frontal *δ* oscillations characteristic of waking^23, 24^ could be used to distinguish epochs exhibiting *δ-*HFO PAC. On the other hand, *θ*-HFO PAC was correlated with CA1 *θ* power, increased during waking locomotion in all conditions, and was observed only during REM epochs exhibiting an elevated ratio of CA1 *θ* power to frontal *δ* power.

### 3.1 HFO Phase Modulation and Control of Mesolimbic Circuits

Multiple pieces of evidence indicate that the HFOs potentiated by NMDAR antagonism and recorded throughout the brain are associated with mesolimbic circuits^15–19^, which receive inputs from both frontal cortex and limbic regions including the hippocampus. HFO power and coupling are modulated not only by ketamine, but also by sleep state^39, 40^, serotonergic^41^ and DAergic^10, 42, 43^ drugs, the level of cognitive control required by a serial reaction time task^44^, the reward probability of a probabilistic reinforcement learning task^45^, and fear conditioning and extinction^15, 17^. While their physiological significance remains unknown, these findings suggest that HFOs play a role in the context-and state-dependent deployment of memory and rule-based strategies for goal-directed behavior, functions in which cortico-hippocampo-mesolimbic circuits are implicated.

Given that HFOs reflect the population activity of mesolimbic structures^15–19^, patterns of HFO PAC may primarily report the dynamic states of these networks. Our results indicate that *θ*-HFO coupling may be a signature of functional connectivity between hippocampal and mesolimbic networks, while *δ*-HFO coupling signifies cortico-mesolimbic connectivity. Recent research shows that in frontal-basal ganglia networks, dopamine agonism can switch the frequency of phase modulation of *γ* and HFO amplitude, from *δ* to *θ*^43, 46^. We suggest that one possible mechanism mediating this switch – at least in the case of HFOs – may be the potentiation of hippocampo-accumbal synapses^25–27,29^. Similarly, the reported state dependence of *θ*-HFO coupling^39^, which we here show relies on the concurrent expression of a high ratio of CA1 *θ* to frontal *δ* power, may reflect the predominance of hippocampo-mesolimbic connectivity during REM.

Notably, recent research on respiratory-related oscillations (RROs) has found that widespread endogenous cortical rhythmic activity synchronized with the respiratory rhythm occurs at both *δ* and *θ* frequencies^47–50^. These RROs may phase-modulate higher frequency rhythms including *γ* and HFOs, and transitions between *δ*-*γ* PAC and *θ*-*γ* PAC can reflect changes in the respiratory frequency during inactive vs. active behavioral states, respectively^51^. It seems unlikely that this phenomenon explains our data, or other data on HFO phase modulation, as such shifts in coupling with the respiratory frequency have not been observed for HFOs^51^, and in our data changes in HFO PAC occurred even within active wake epochs. However, we did not record respiration, and it is possible that the changes between *δ*-HFO and *θ*-HFO PAC observed here align with simultaneous changes in respiratory frequency.

Behaviorally, hippocampo-accumbal inputs and D1 receptor activation act in concert to enable focus on the current task and execution of the current strategy or action, while cortico-accumbal inputs and D2 receptor inactivation enable flexible switching of focus and strategy^52, 53^. One primary role of the mesolimbic system, and the NAcc in particular, may be to control the switch between flexible and context-driven strategies in goal-directed behavior, something that may be accomplished in part through the selection of cortical vs. hippocampal inputs^52, 53^. Our results suggest that HFO phase-coupling tracks and may even mediate this input selection. Alterations in HFO phase-coupling seen with pharmacological interventions may thus play a key role in the changes in executive function and goal-directed behavior observed with these interventions.

Examining both *θ* and *δ* modulation of HFO amplitude is crucial for understanding the functional significance of HFO PAC. In a recent study of human performance on a serial reaction time task^44^, NAcc *θ*-HFO PAC correlated with reaction times, correlated inversely with error rates, and was elevated during task blocks requiring subjects to access newly learned stimulus-response associations (“HCC blocks”) relative to task blocks during which subjects repeated a learned motor pattern (“LCC blocks”). These data support the hypothesis that *θ*-HFO coupling mediates greater focus on task execution by increasing hippocampo-accumbal connectivity. However, the authors speculated that *θ*-HFO PAC signaled “a deviation from expectancy” implying “the need to stop an automated motor routine” executed during LCC blocks^44^. In contrast, we suggest that LCC blocks represented an HFO PAC-free low-engagement state of habitual responding, while HCC blocks represented a high engagement state of focused task execution. While the authors examined HFO PAC only at phase frequencies above 3 Hz, we suggest that *δ-*HFO PAC, representing a high engagement state of cortico-mesolimbic drive and strategy re-evaluation, should be observable briefly after shifts from LCC to HCC blocks.

### 3.2 Subunit-Specific Oscillopathies Dissociate Frontal Disinhibition and Hippocampal Hyperactivity

A large body of research implicates an altered balance between prefrontal and hippocampal activity^11, 54–56^ – and its downstream effects on affect, reward, and motivation circuitry^52, 57, 58^ – in the pathophysiology of schizophrenia. It has been hypothesized that the selective susceptibility of PV+ inhibitory interneurons to NMDAR blockade may result in the disinhibition of PFC pyramidal cells, and a subsequent increase in “baseline” activity and *γ* power^11, 59, 60^ – a paradoxical case of hyperactivity leading to hypofunction^61^. The increases in high *γ* and HFO power and in *δ*-HFO PAC seen in this study with both nonspecific and NR2A-preferring blockade suggest an overlap between the mechanisms generating increased cortical *γ* power^14^ and those generating increased HFO activity. In contrast, the signatures of hippocampal hyperactivity we observed following nonspecific NMDAR blockade – increased *θ*-HFO PAC and increased hippocampal *θ*-*γ* PAC – were absent following NR2Apreferring NMDAR blockade. Indeed, NVP-AAM077 induced a marked decrease in CA1 *θ* power, suggesting a suppression of hippocampal function. Thus, disinhibition of PFC and hippocampal hyperactivity appear to be dissociable via subunit-specific NMDAR blockade.

We hypothesize that the abnormal increases in *δ*-HFO PAC and HFO power observed following NVP-AAM077 administration are signatures of prefrontal disinhibition, induced via blockade of NMDARs of NR2A subtype preferentially expressed by PV+ interneurons. However, the blockade of NMDARs in nucleus reuniens of the thalamus, an important node mediating interactions between PFC and hippocampus^24, 62^, also leads to altered prefrontal-hippocampal balance as well as to overexpression of *δ* rhythms in thalamus and limbic structures^63–67^. Additionally, the relationship between the narrowband *δ* described here and previously^24^ and the respiratory rhythm^47–51, 68^ is an important question for further investigation.

It is tempting to hypothesize that the distinct HFO coupling profile observed following NR2A-preferring blockade may contribute to the pathological information processing seen with NVP-AAM077 and MK-801, but not Ro25, administration, through dysfunction of executive control mechanisms. Indeed, our results suggest that the increase in *δ*-HFO PAC observed following MK-801 injection is not merely a “spectral leakage” artifact of increased *θ*-HFO PAC, but rather that both *θ*-HFO and *δ*-HFO coupling are potentiated by nonspecific NMDAR antagonism, although the expression of increased *δ*-HFO PAC seems to depend on the concurrent presence of potentiated *θ*-HFO PAC. Interactions between hippocampal and prefrontal NAcc afferents are abnormal in a developmental model of schizophrenia^52, 58^, and an increase in both hippocampo-and cortico-mesolimbic drives, resulting in “focused attention on multiple contingencies”, may play a role in the “disrupted focus and … overwhelming bombardment of stimuli” of schizophrenia^52^. Increased PFC drive to motivational circuits is also hypothesized to be a causal factor in depression^69^, and blockade of NR2A-subtype NMDARs may contribute to the acute induction of negative symptoms with NMDAR antagonism. Further exploration of the behavioral effects of subtype-specific NMDAR antagonism, and their relationship to the altered patterns of rhythmic activity reported here, will be crucial to testing these hypotheses.

## Acknowledgements

This study was supported in part by NSF grant DMS-1042134 (BRPP) and NIH grants HL102241 (KH), AG048108 (KH), MH100820 (BK), and ES006189 (BK). The NVP-AAM077 compound was provided by Dr. Yves P. Auberson from Novartis Institute of Biomedical Research, Basel, Switzerland. We thank two anonymous reviewers for their helpful comments.

## Author contributions statement

BK conceived and performed the experiments; BRPP, BK, and KH conceived the data analysis; BRPP performed the analyses. BRPP drafted the manuscript and figures; all authors reviewed the manuscript.

## Additional information

**Competing interests:** the authors declare they have no competing interests.

## Supplementary Information

**Figure S1.**
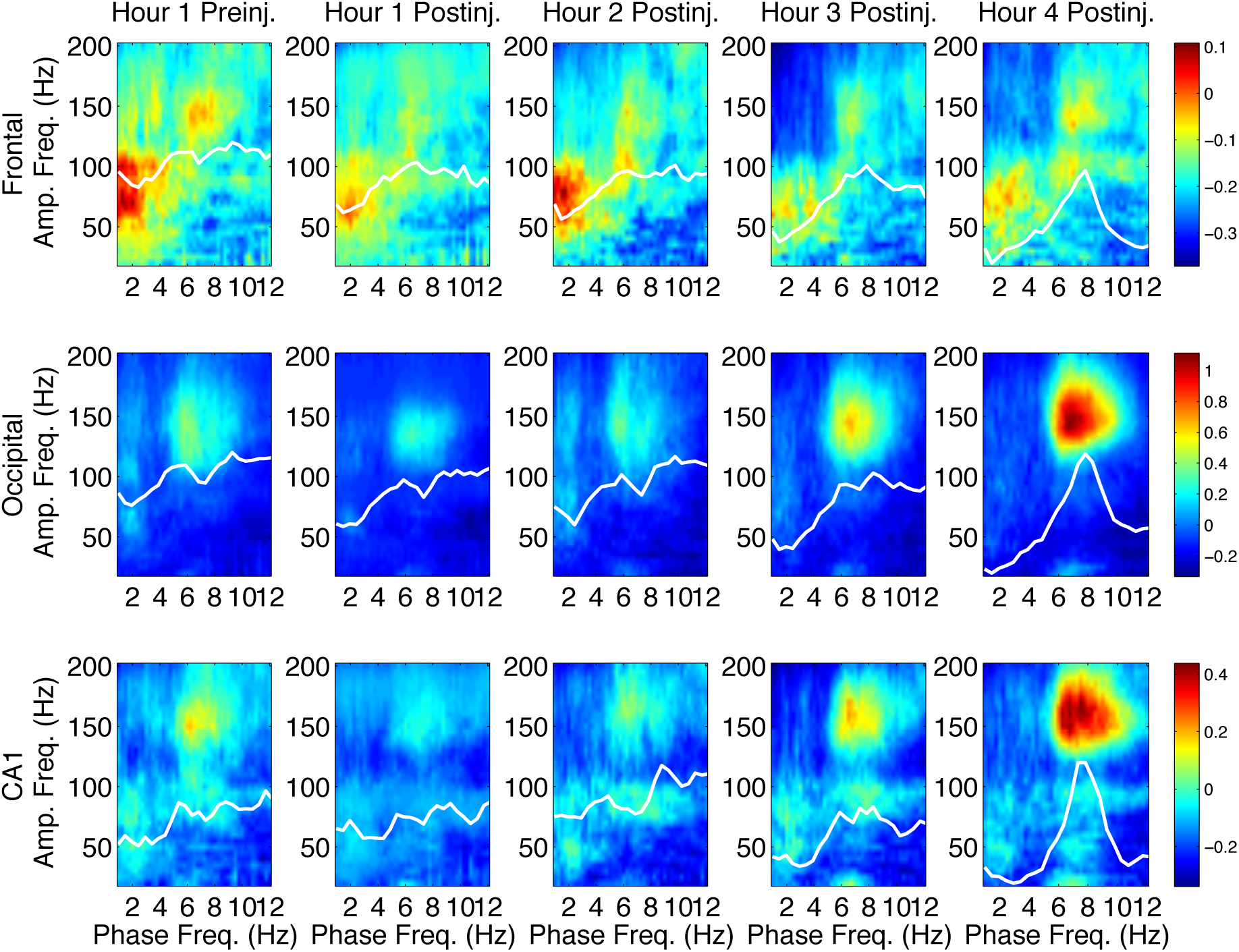
Region-specific PAC patterns following saline injection. Median (n=6) comodulograms for the 1 h preceding and the 4 h following saline injection, shown for frontal (top), occipital (middle), and CA1 (bottom) electrodes (simultaneous recordings). Overlaid white traces show the mean spectral power at frequencies below 12 Hz for each h.

**Figure S2.**
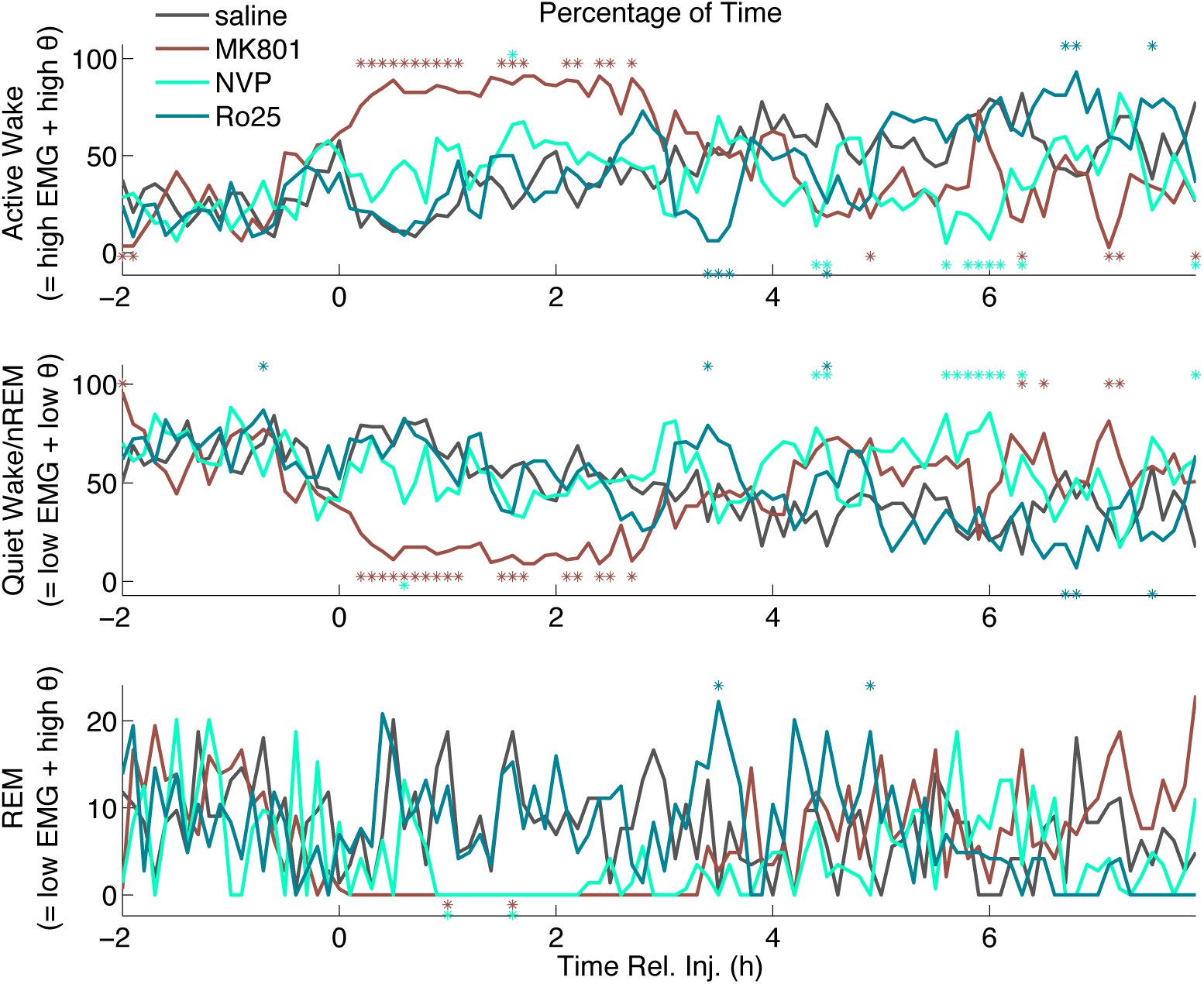
Vigilance states following drug administration. Median (n=6) percentage of each 6 min period spent in AW (defined as high EMG and *θ* power; top), QW/nREM sleep (defined as low EMG and *θ* power, and including quiet waking as well as slow-wave sleep periods; middle), and REM (defined as atonia and high *θ* power; bottom), following injection with saline, MK-801, NVP-AAM077, and Ro25–6985. Stars indicate 6 min periods for which spectral power is significantly higher (above) or lower (below) than saline (ranksum test, significance level = 0.05).

**Figure S3.**
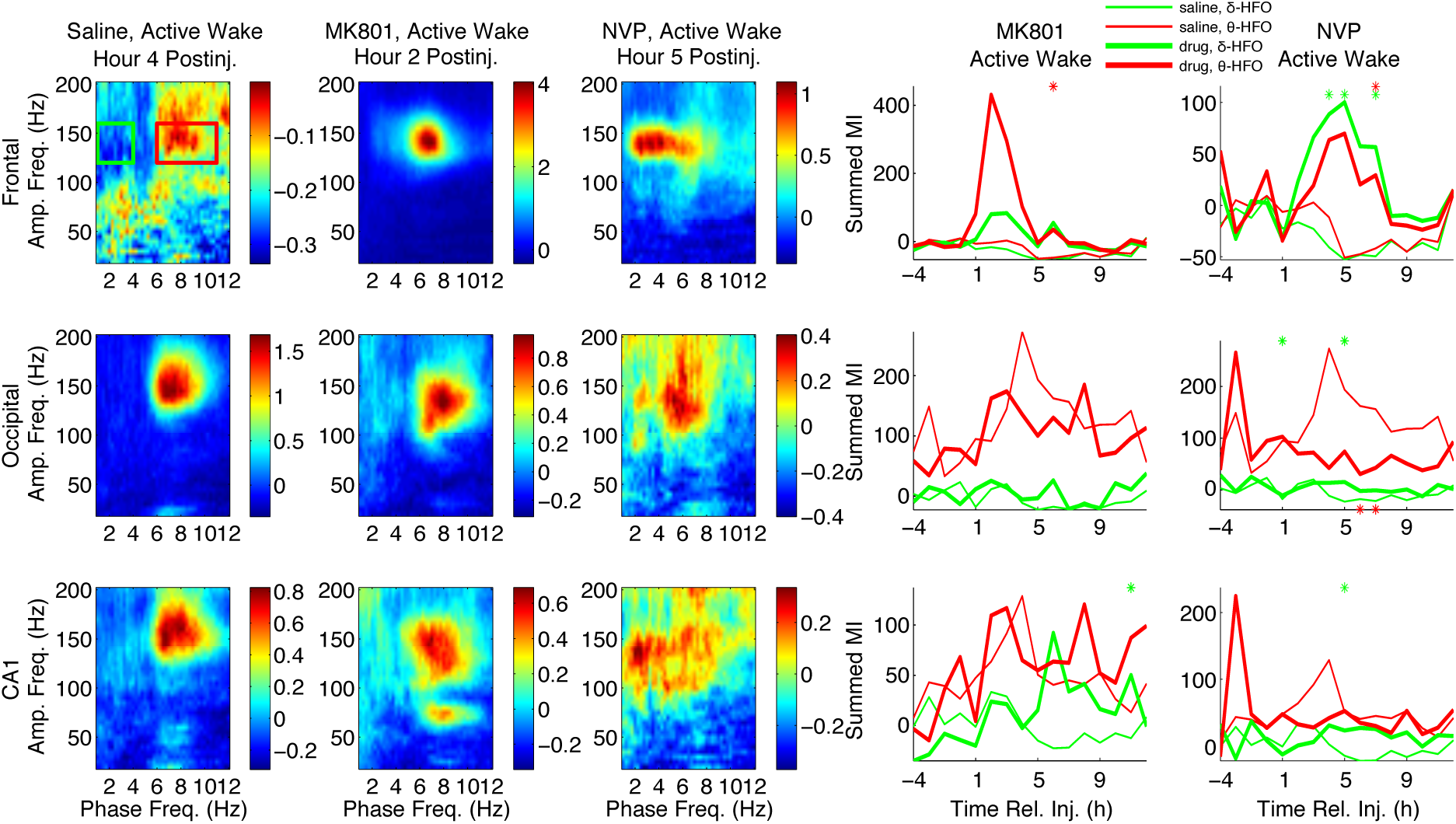
Patterns of phase-amplitude coupling following MK-801 and NVP-AAM077 injection differ markedly during AW epochs. Left three columns show median (n=6) comodulograms for all AW epochs during the h of largest effect for simultaneous LFP recordings from frontal (top), occipital (middle), and CA1 (bottom) electrodes, following injection of saline, MK-801, and NVP-AAM077. Right two columns show timeseries of summed PAC over the *θ*-HFO (red rectangle and time series) and *δ-*HFO (green rectangle and time series) frequency ranges for all AW epochs from 4 h pre-injection to 12 h post-injection. Thick lines are drug time series; thin lines are saline time series; stars indicate 6 min periods for which drug PAC is significantly higher (above) or lower (below) than saline (ranksum test, significance level = 0.05).

**Figure S4.**
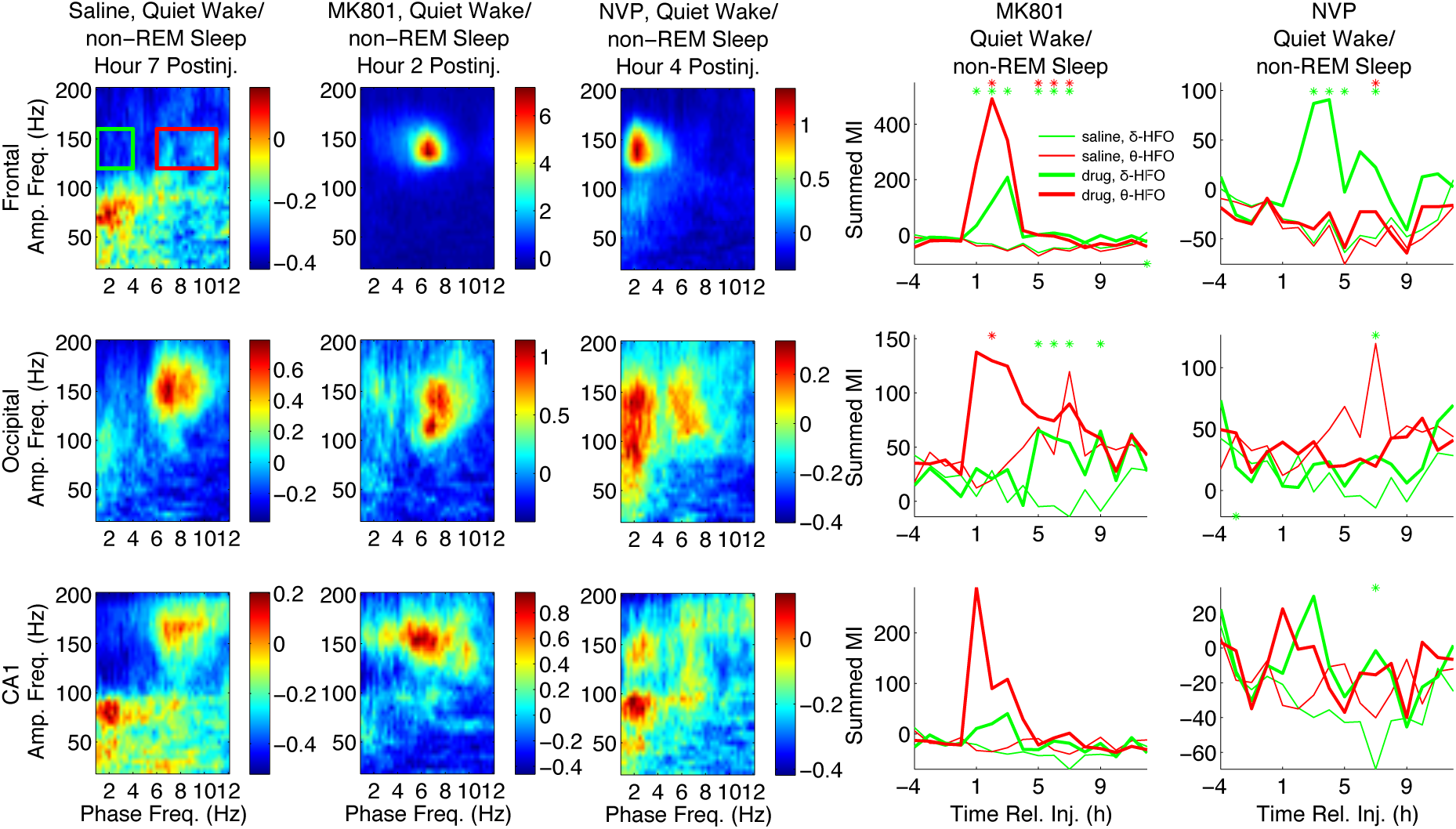
Patterns of phase-amplitude coupling following MK-801 and NVP-AAM077 injection differ markedly during QW/nREM epochs. Left three columns show median (n=6) comodulograms for all QW/nREM epochs during the h of largest effect for simultaneous LFP recordings from frontal (top), occipital (middle), and CA1 (bottom) electrodes, following injection of saline, MK-801, and NVP-AAM077. Right two columns show timeseries of summed PAC over the *θ-*HFO (red rectangle and time series) and *δ-*HFO (green rectangle and time series) frequency ranges for all QW/nREM epochs from 4 h pre-injection to 12 h post-injection. Thick lines are drug time series; thin lines are saline time series; stars indicate 6 min periods for which drug PAC is significantly higher (above) or lower (below) than saline (ranksum test, significance level = 0.05).

**Figure S5.**
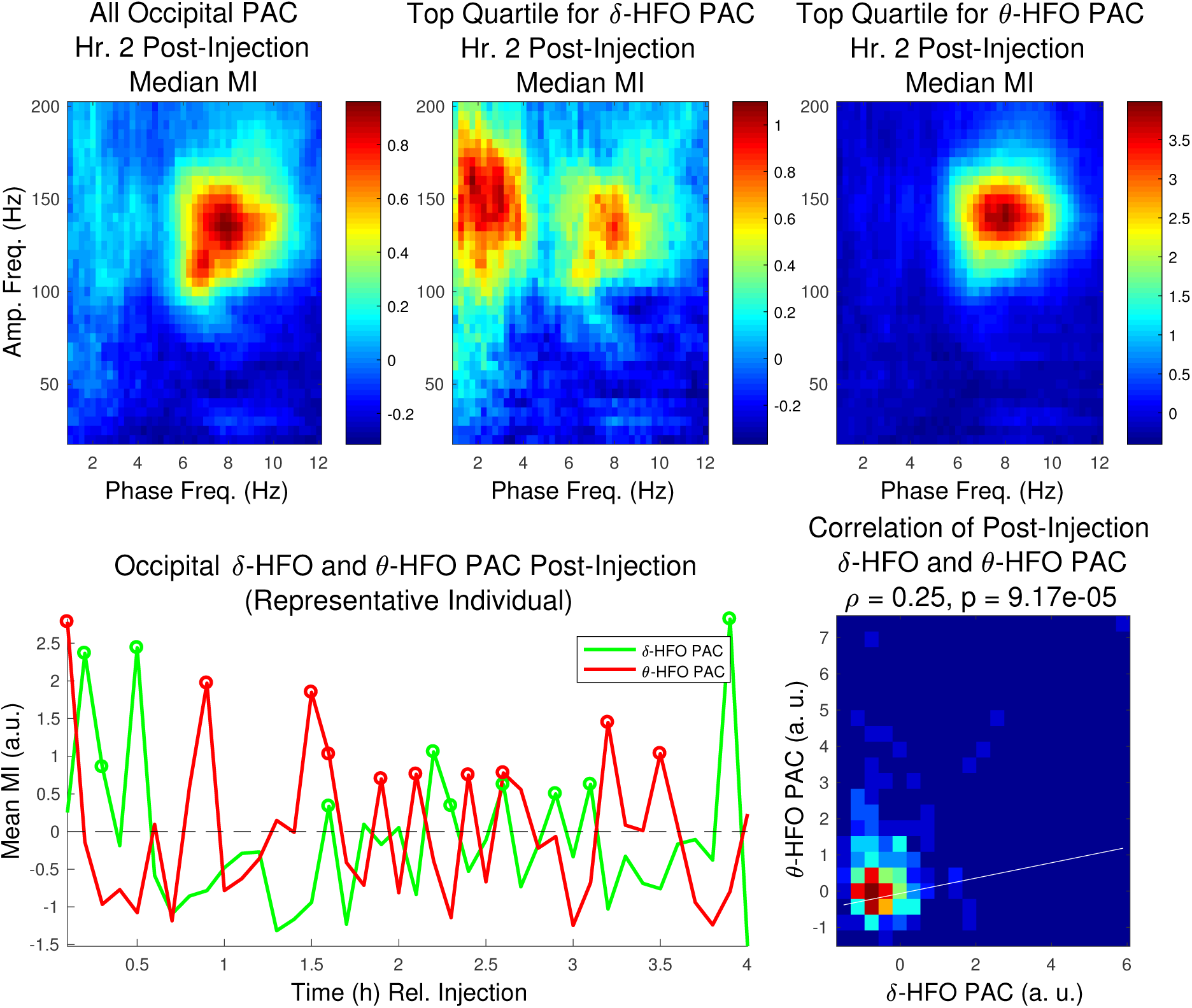
*δ*-HFO PAC and *θ*-HFO PAC are correlated in occipital cortex following MK-801 administration. Top left, median occipital comodulograms in the 2*^nd^* h following MK-801 injection. Top middle and right, median occipital comodulograms for the top quartile of epochs with respect to *δ-*HFO PAC and *θ-*HFO PAC. Bottom left, time courses of summed *δ-*and *θ-*HFO PAC for 6 min. periods following injection (representative animal, circles indicate 6 min. periods in the top quartile). Bottom right, the correlation between *δ-*and *θ-*HFO PAC over all animals.

**Figure S6.**
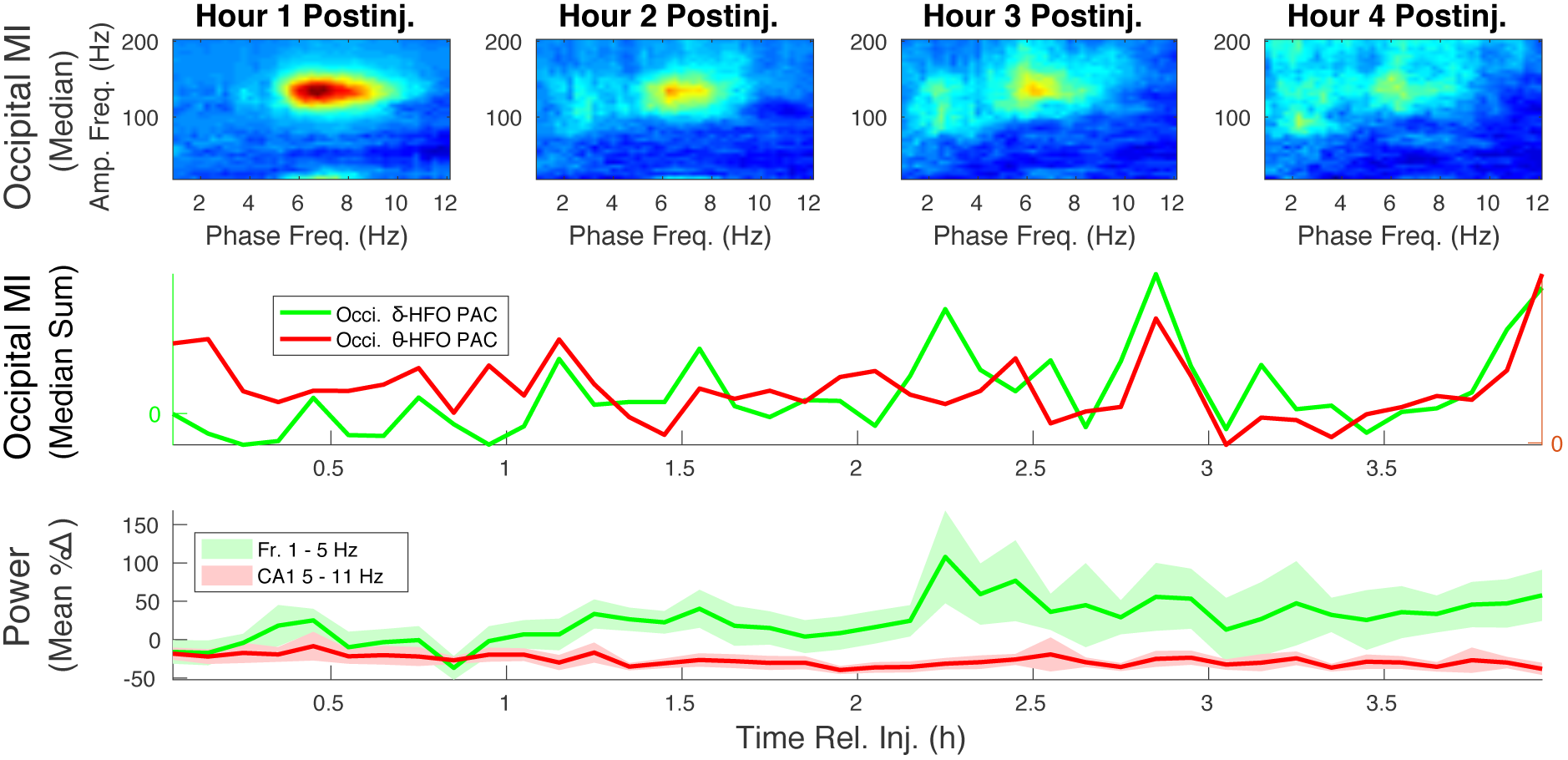
The time courses of frontal *δ* power and CA1 *θ* power mirror the time courses of *δ-*HFO PAC and *θ-*HFO PAC, respectively, during AW epochs. Top, median (n=6) occipital comodulograms are shown for AW epochs during the first 4 h following NVP-AAM077 injection. Middle, profiles of median (n=6) summed PAC during AW epochs (at 6 min intervals) are shown for *δ-*HFO and *θ*-HFO ranges. Bottom, mean ± s.d. (n=6) frontal *δ* power (summed over 1–5 Hz) is plotted alongside median (n=6) CA1 *θ* power (summed over 5–11 Hz) for AW epochs.

**Figure S7.**
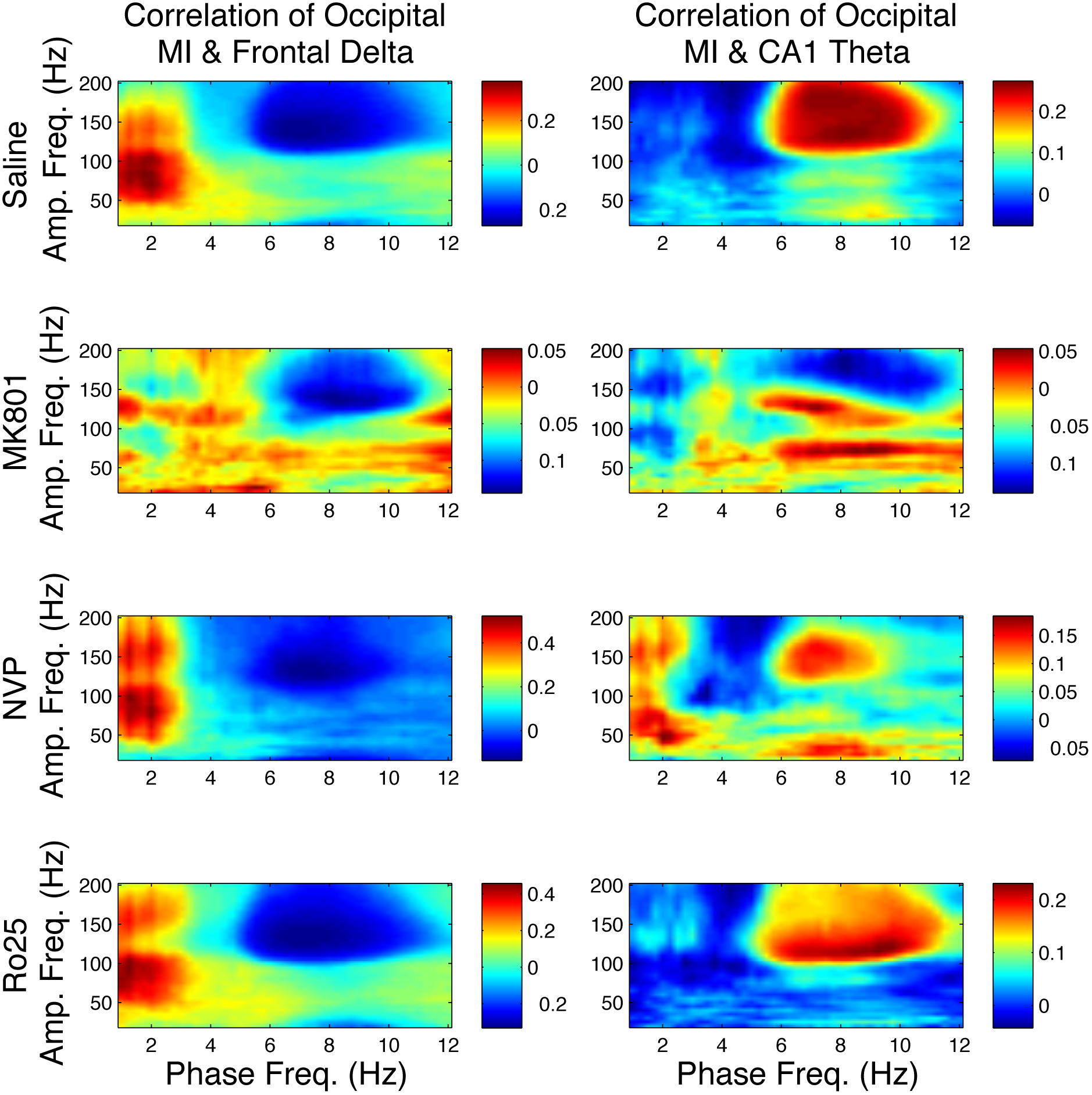
Frontal *δ* and CA1 *θ* are correlated with *δ-*HFO and *θ-*HFO PAC. The correlation between PAC MI and Frontal *δ* power (left) or CA1 *θ* power (right) during the first 4 h following injection are shown for each pair of phase-giving and amplitude-giving frequencies, and for each channel, frontal (top), occipital (middle), and CA1 (bottom).

**Figure S8.**
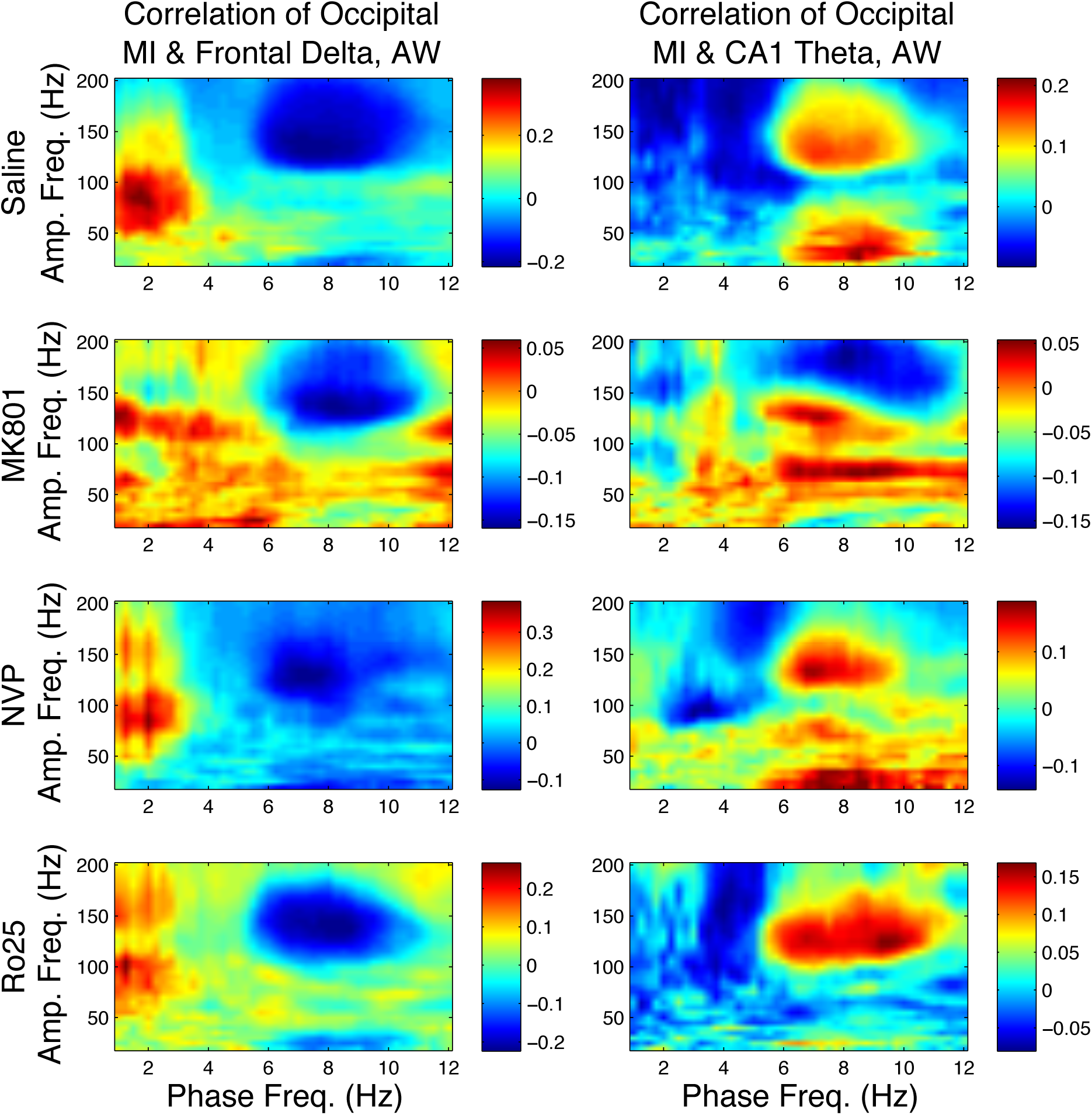
Frontal *δ* and CA1 *θ* are correlated with *δ-*HFO and *θ*-HFO PAC during AW epochs. The correlation between PAC MI and Frontal *δ* power (left) or CA1 *θ* power (right) during AW epochs for the first 4 h following injection are shown for each pair of phase-giving and amplitude-giving frequencies, and for each channel, frontal (top), occipital (middle), and CA1 (bottom).

